# Guiding the design of well-powered Hi-C experiments to detect differential loops

**DOI:** 10.1101/2023.03.15.532762

**Authors:** Sarah M Parker, Eric S. Davis, Douglas H. Phanstiel

## Abstract

3D chromatin structure plays an important role in gene regulation by connecting regulatory regions and gene promoters. The ability to detect the formation and loss of these loops in various cell types and conditions provides valuable information on the mechanisms driving these cell states and is critical for understanding how long-range gene regulation works. Hi-C is a powerful technique used to characterize three-dimensional chromatin structure; however, Hi-C can quickly become a costly and labor-intensive endeavor, and proper planning is required to determine how to best use time and resources while maintaining experimental rigor and well-powered results. To facilitate better planning and interpretation of Hi-C experiments, we have conducted a detailed evaluation of statistical power using publicly available Hi-C datasets paying particular attention to the impact of loop size on Hi-C contacts and fold change compression. In addition, we have developed Hi-C Poweraid, a publicly-hosted web application to investigate these findings (http://phanstiel-lab.med.unc.edu/poweraid/). For experiments involving well-replicated cell lines, we recommend a total sequencing depth of at least 6 billion contacts per condition, split between at least 2 replicates in order to achieve the power to detect the majority of differential loops. For experiments with higher variation, more replicates and deeper sequencing depths are required. Exact values and recommendations for specific cases can be determined through the use of Hi-C Poweraid. This tool simplifies the complexities behind calculating power for Hi-C data and will provide useful information on the amount of well-powered loops an experiment will be able to detect given a specific set of experimental parameters, such as sequencing depth, replicates, and the sizes of the loops of interest. This will allow for more efficient use of time and resources and more accurate interpretation of experimental results.

**HIGHLIGHTS:**

1. Distance-dependent Hi-C contacts and fold change compression influence power
2. Integrative modeling of Hi-C power provides guidance for experimental design
3. An interactive web app aids the design of differential Hi-C experiments

## INTRODUCTION

3D chromatin structure is thought to play a critical role in the regulation of gene expression, particularly during development and in response to external stimuli^1–3^. Abnormalities in this organization have been implicated in a number of human diseases and developmental disorders^4^. While multiple types of 3D chromatin structures have been identified, chromatin loops—point-to-point interactions between two genomically distal loci—are of particular interest as they can bring gene promoters into close physical proximity with distal enhancers and facilitate transcriptional activation^5^. Multiple genomic approaches have been developed to detect loops and other chromatin structures, including Hi-C, a widely used approach to quantify chromatin interactions in a genome-wide fashion^6–9^. Computational analysis of Hi-C data can be used to both identify chromatin loops and quantify the interaction frequencies between pairs of loop anchors. The application of Hi-C to investigate differential looping, or changes in looping across samples or biological conditions, has provided valuable insights into the function of the human genome, the role of enhancers in regulating gene expression, and the genetic basis of human disease^2,10^.

Proper design and interpretation of comparative Hi-C genomics experiments requires a rigorous understanding of the statistical power underlying the experiment in question. Statistical power is the probability of a test rejecting the null hypothesis, given that the null hypothesis is false. In this case, the null hypothesis is that a given loop is not changing between conditions. Power relies on several key factors including the sample size (e.g. biological replicates), the effect size (i.e. fold change), the counts representing the feature of interest (i.e. the loop pixel), the alpha level, and dispersion. Several software packages have been developed to estimate the power of comparative genomic studies and these tools have been extensively used to design and interpret studies involving RNA-seq, ChIP-seq, ATAC-seq, and numerous other genomic methodologies^11–15^. However, Hi-C has several inherent differences compared to these data types that make power analysis non-trivial. First, the generation of Hi-C data sets for preliminary power analysis is expensive since Hi-C requires sequencing depths that are orders of magnitude greater than those required for traditional sequencing experiments. Second, Hi-C library preparation and data analysis both require special expertise due to the lengthy protocol and the sheer size and complexity of the resulting data sets. Finally, the counts observed in a given Hi-C pixel arise due to multiple biological and technical forces, each of which is dependent on the genomic distance between the loci depicted by said pixel.

Hi-C interaction frequencies are governed by at least two main forces, or types of interactions, each of which affects the power to detect differential looping. It is the cumulative influence of these interaction types that gives rise to the counts observed at any particular pixel in a Hi-C dataset. The first type of interactions are **polymer interactions**, which are distance-dependent interactions between two regions of a chromosome due to their inclusion in a linear polymer. The second type are **looping interactions**, interactions driven by the forces of specific chromatin interacting proteins. Most looping interactions are presumably the result of CTCF binding and cohesin-driven loop extrusion; however, other less common mechanisms have also been identified^16–18^. Other forces, including compartmentalization and contact domain inclusion, also influence interaction frequencies albeit to a lesser degree. An understanding of the relationships between these forces can strongly influence the power to detect differential loops as we describe here.

To facilitate better planning of Hi-C experiments, we have modeled these forces and conducted a detailed evaluation of the effects of sequencing depth and replicate number on differential loop detection using deeply sequenced, publicly available Hi-C data sets^19^. We also developed an interactive web application to enable better planning of comparative Hi-C experiments. Finally, we provide guidance and recommendations for planning Hi-C experiments depending on project goals and budgets.

## RESULTS

### Loop size is anti-correlated with sequencing counts

One of the key determinants of power in genomic experiments is the number of sequencing counts attributed to the feature being measured, in this case, loops. One of the unique aspects of Hi-C data is that on average, sequencing counts decrease as a function of genomic distance. We modeled this trend to understand how it affects the power to detect differential loops at various distances. Using deeply sequenced Hi-C data from Rao & Huntley et al., which includes roughly 5 billion unique contacts in GM12878 cells, we extracted the observed counts and expected counts for all loops and plotted the median counts as well as interquartile and interdecile ranges of observed counts (**Fig 1A**). These counts decrease as genomic distance increases, and the relationship follows a power law curve as previously observed^6^. This suggests that the size of a differential loop is closely correlated with the power we have to detect it. To elucidate this relationship, we calculated the power to detect differential loops (via the RNAseqpower package^11^) using the median counts for each genomic distance as values for depth (**Fig 1B**). We held dispersion, alpha, and fold change constant at 0.001, 0.05, and 2, respectively. As expected, power decreases sharply with increasing loop size, an effect that is slightly alleviated with increased sequencing replicates. The distribution of loop sizes is skewed toward the shorter range of the distances shown (**Fig 1C**). These trends were similar for other Hi-C data sets investigated(**Fig S1**)^1^. How the distributions of loop sizes and power intersect to affect the overall power of a differential Hi-C experiment will be explored in more detail later. Examples of loops at varying distances and their associated statistical power are depicted in **Figure 1D**.

**Figure 1.**
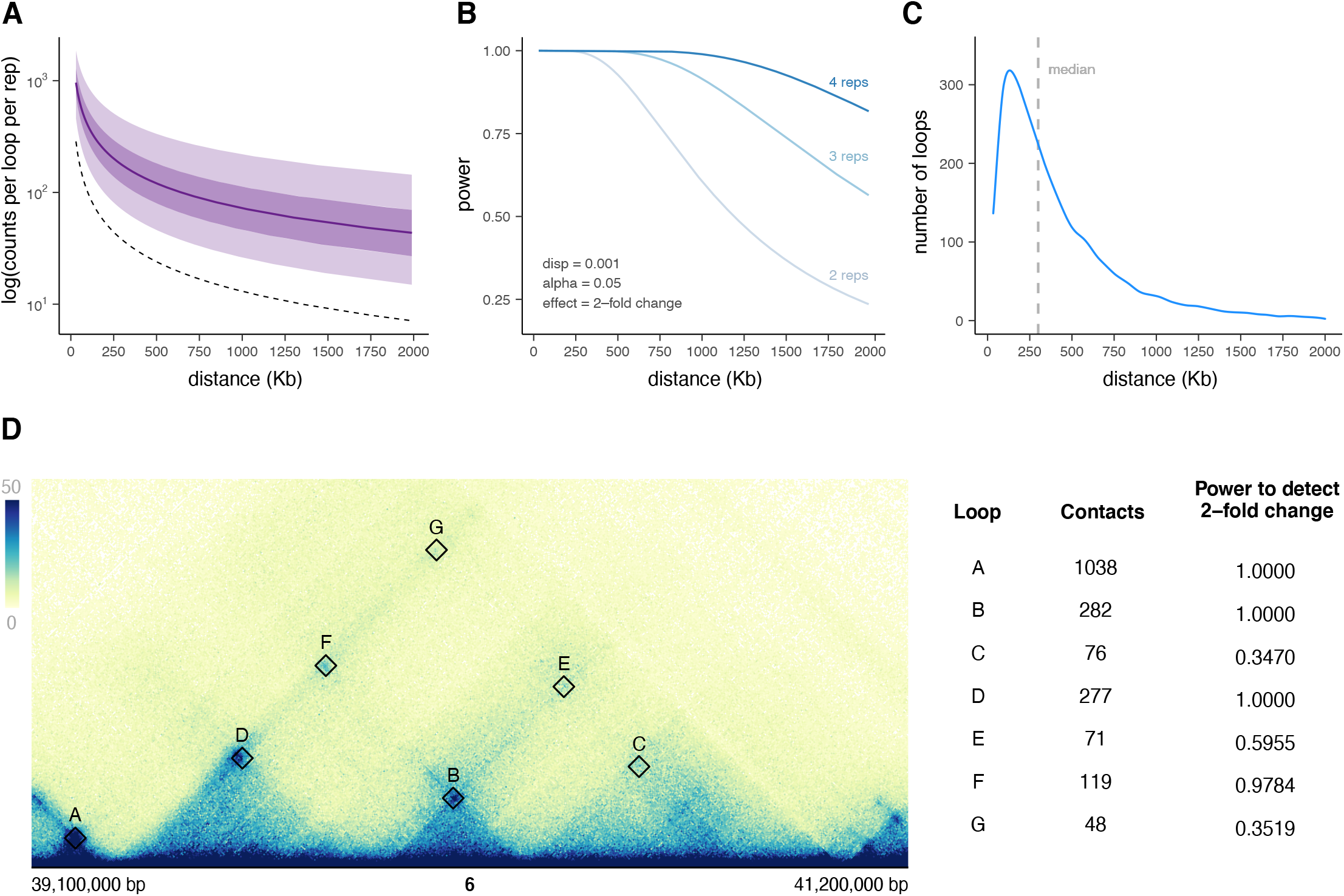
Counts and power decrease with increasing loop size. **(A)** Median counts per loop as a function of genomic distance are plotted as a solid purple line. Dark and light purple shaded areas represent inner interquartile and interdecile ranges. The dotted black line represents the median value of all (loops and non-looped) pixels at each 10 Kb binned genomic distance. **(B)** Effect of loop size on power when dispersion, alpha, and effect are held constant at 0.001, 0.05, and 2, respectively. This distance-dependent effect is due only to changes in counts and is investigated across different replicate values. **(C)** Distribution of loop sizes showing a median loop size of 300 Kb. **(D)** An example region on chromosome 6 with loops labeled in order of increasing distance. The observed contacts and the power to detect a 2-fold change in counts due to looping interactions are listed for each loop pixel.

### Loop size is anti-correlated with fold change compression

A second key determinant of statistical power is the effect size—or fold change—of the features of interest. However, since only a fraction of the counts at a given loop pixel arise from looping interactions, observed fold changes in a Hi-C experiment are smaller than the actual changes in chromatin looping. To illustrate this compression, we consider a single 350 Kb loop from the GM12878 data set acquired by Rao & Huntley et al^19^ **(Fig 2A)**. There are 159 observed counts at this loop pixel. Polymer interactions for this pixel can be estimated by calculating the median inter-actions for all pixels connecting loci 350 Kb apart. Because the vast majority of pixels do not represent a chromatin loop, the resulting value is called the ‘expected’ counts since this is the number of counts that would be expected in the absence of a chromatin loop. For the loop in question, this provides a value of 36 counts, or 23% of the observed counts. Looping interactions can be estimated by subtracting the expected counts from the observed counts which gives us 123, or 77% of the observed counts. Therefore, a two-fold increase in looping interactions (123 * 2 = 246) would actually only be observed as a 1.77-fold increase in observed counts **(Fig 2A)** since the expected counts would remain the same.

**Figure 2.**
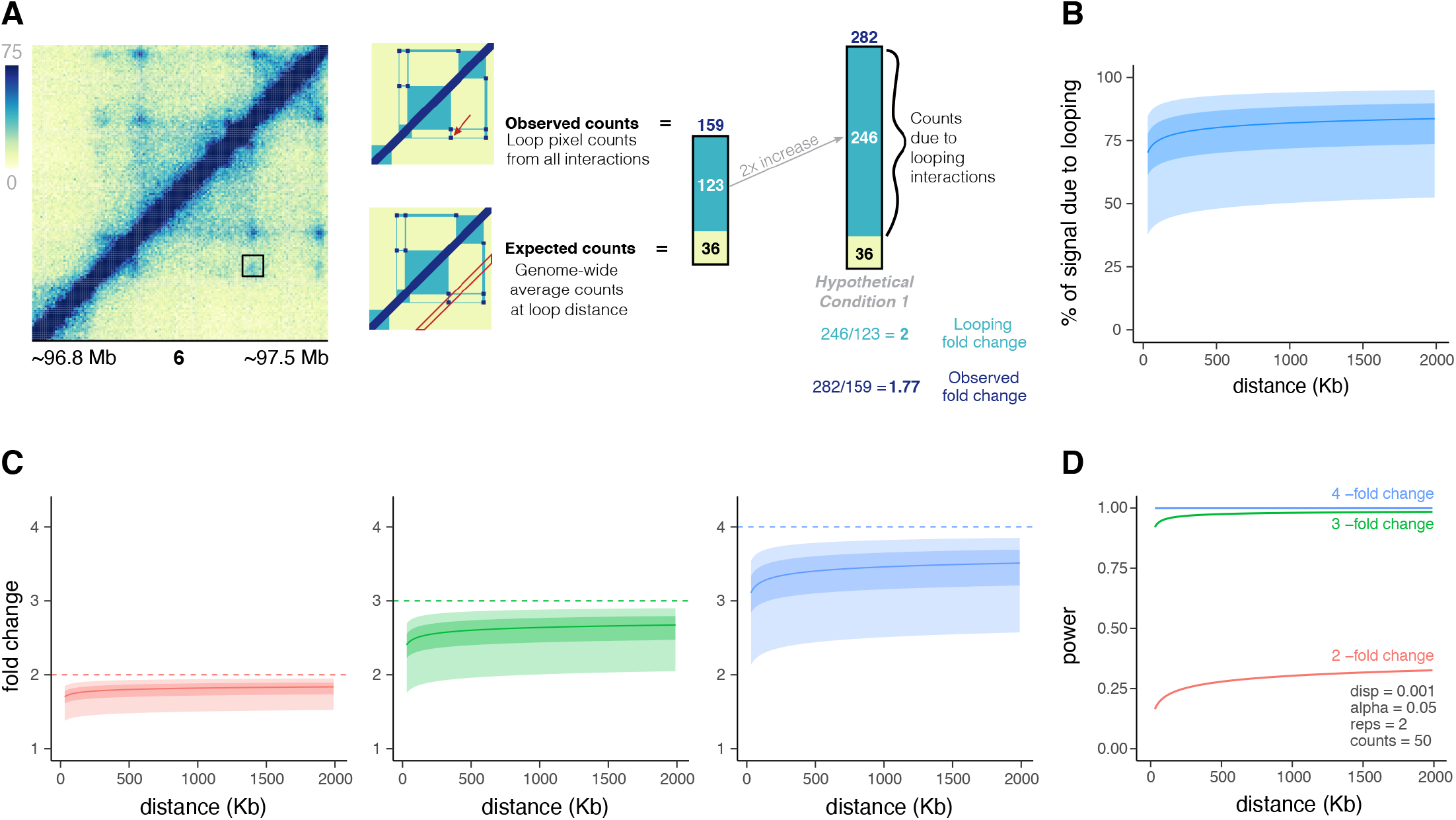
Fold change compression is anti-correlated with loop size. **(A)** An example region showing a loop and the effect on observed counts and fold change if counts due to looping interactions are doubled. The counts due to looping interactions can be represented as the observed counts minus the expected counts. When these looping counts are doubled, the expected counts remain the same, so the observed fold change of total counts (1.77) is less than that of the fold change of looping counts (2)). **(B)** The distance-dependent nature of the percent of signal due to looping. The percent of signal due to looping is about 50-80% of the observed counts and increases slightly with loop size. For smaller loop sizes, expected counts comprise more of the observed counts. **(C)** Observed fold change per loop for looping fold changes of 2 (red), 3 (green), and 4 (blue). Dark and light shaded areas represent interquartile and interdecile ranges. Observed fold change is compressed when looping doubles, triples, and quadruples, with an increasing range of effect as fold change increases. The compression effect is greatest for shorter-range loops, meaning longer-range loops typically have higher observed fold changes for the same change in looping counts. **(D)** Effect of loop size on power for a median loop at each distance when dispersion, alpha, replicates, and counts are held constant at 0.001, 0.05, 2, and 50, respectively.

We next sought to explore this effect across all loops identified in a given Hi-C experiment. Using the same approach described above, we estimated the percentage of counts due to looping interactions for all loops (**Fig 2B**). As loop size increases, we observe a higher percentage of counts due to looping. We next explored how this distance-dependent change in the composition of sequencing counts impacts fold change compression. For every loop, we calculated the percentage of counts due to looping and calculated what the observed fold change would be given a two, three, or four-fold change in looping. The median as well as interquartile and interdecile ranges are plotted in **Figure 2C**. As expected, fold change compression was observed for all loop sizes; however, the magnitude of fold change compression decreased with increasing loop size.

To determine how these distance-dependent effects on fold change compression impact statistical power, we calculated the power to detect the median values of observed fold change for each loop size (**Fig 2D**). Dispersion, alpha, replicates, and counts were held constant at 0.001, 0.05, 2, and 50 respectively. Plotting the resulting power reveals the opposite trend that we observed when considering counts as a function of distance. That is, the correlation between sequencing counts and loop size suggests a positive correlation between loop size and power, whereas the inverse correlation between fold change compression and loop size suggests a negative correlation between loop size and power. However, these models were built by isolating individual variables and using only median values of sequencing counts and fold change per loop size bin. How these features intersect to determine power on a per-loop basis in real data sets and how these relationships impact experimental design is explored below.

### Recommendations for maximizing power

To determine optimal experimental parameters, such as sequencing depth and replicates, we calculated power for every loop in the deeply sequenced dataset created by Rao & Huntley et. al^19^. We subsampled the dataset to 10 different sequencing depths and calculated power using a range of both dispersions and replicates. Observed fold changes were modeled using the fold change compression relationships for each loop as calculated in **Figure 2.** For the case of this analysis, a well-powered loop is defined as one which has 80% or higher power to detect a 2-fold change in looping. As expected, a higher overall sequencing depth results in increased power. At low values of dispersion, (those we typically observe when replicates are defined as different wells of the same cell line) the number of replicates does not heavily influence power. The total sequencing depth is the main determinant of power, and the number of well-powered loops is similar if those reads are split across two or ten replicates. For experiments with higher dispersion, (for example, if replicates come from donors with different genetic backgrounds, ages, ethnicity, etc) increasing replicates heavily influences power, with more replicates always being better. Even at relatively high sequencing depths, (i.e. 6 billion contacts per condition) power is highly distance dependent with shorter loops being far more well-powered than longer loops. The percent of well-powered loops at each distance is shown in **Fig 3B-C.** Fortunately, loop size distributions are skewed to shorter loops so the low power to detect loops greater than 1 Mb does not heavily influence the overall power of the experiment. Nevertheless, for proper interpretation of differential Hi-C experiments, it is important to consider this bias toward the detection of shorter loops.

**Figure 3.**
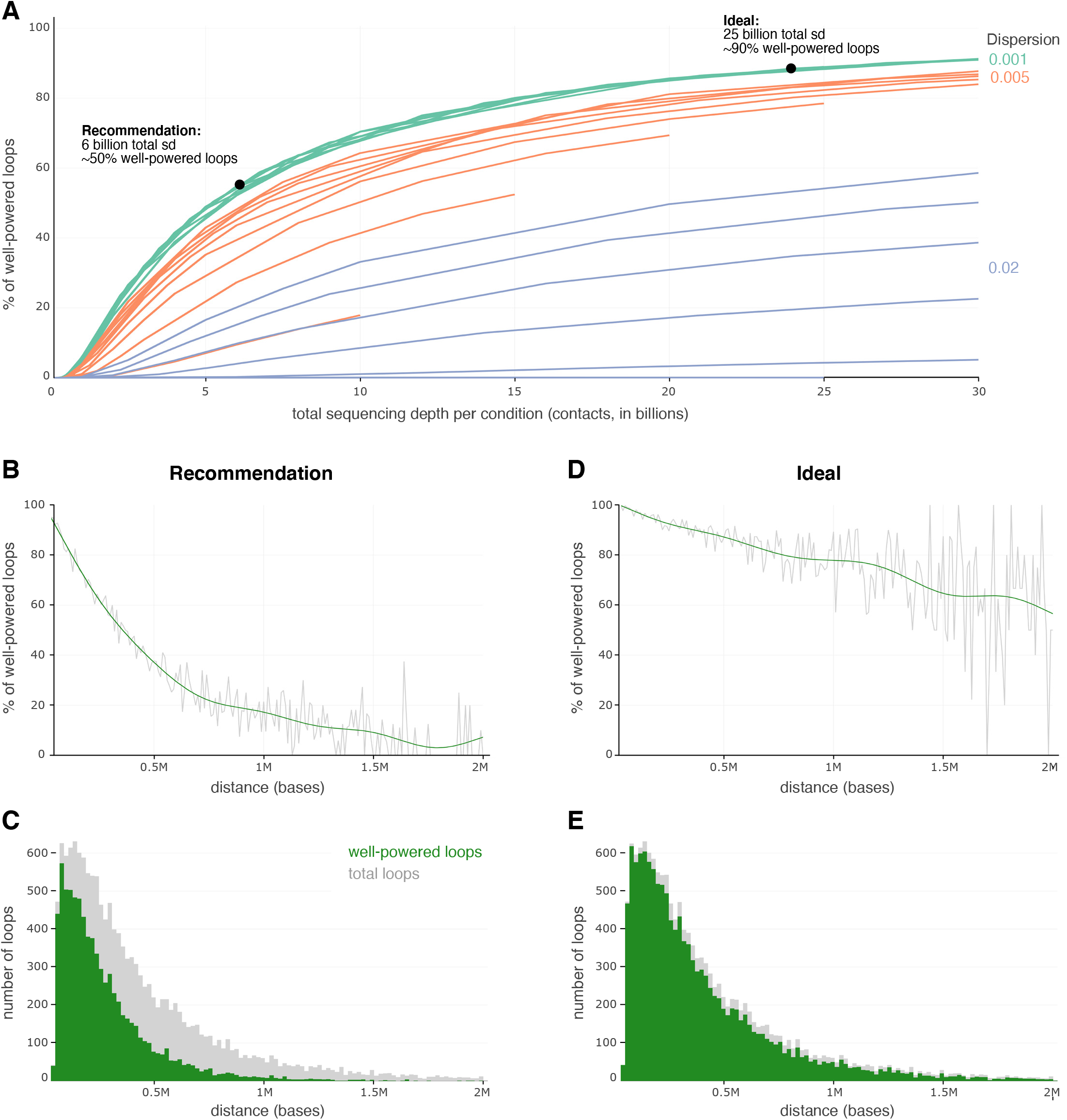
Power increases with replicates and sequencing depth. **(A)** For each combination of replicates (2-10), dispersion (0.001, 0.005, and 0.02), and sequencing depth per replicate (50 M, 100 M, 250 M, 500 M, 750 M, 1 B, 2B, 3 B, 4 B, and 5 B contacts), the power to detect a 2-fold change in looping was calculated. Loops with a power above 0.8 were designated as “well-powered”, and the percentage of these loops is represented on the x axis. This percentage is investigated across the total sequencing depth per condition (multiplying replicates by the sequencing depth per replicate). For the lowest dispersion, we highlight 2 different scenarios: a recommendation of 6 billion contacts per condition to achieve ~50% well-powered loops, and a more ideal scenario of 25 billion contacts per condition to achieve ~90% well-powered loops. **(B–C)** The percent of well-powered loops **(B)** and number of loops **(C)** in Rao & Huntley et al.’s GM12878 data for the recommended **6 billion** contacts per condition, using dispersion and replicate values of 0.001 and 2, respectively. The power to detect a 2-fold change in looping interactions was calculated per loop, and the percentages and numbers of loops with a power over .8 were calculated per 10Kb bin. **(D–E)** The percent of well-powered loops **(D)** and number of loops **(E)** in Rao & Huntley et al.’s GM12878 data for the ideal **25 billion** contacts per condition, using dispersion and replicate values of 0.001 and 2, respectively. The power to detect a 2-fold change in looping interactions was calculated per loop, and the percentages and numbers of loops with a power over .8 were calculated per 10Kb bin.

Based on these findings, we recommend sequencing to a depth of at least 6 billion contacts per condition. This will ensure that roughly 50% of loops will exhibit 80% power to detect a two-fold change in looping. These calculations assume that replicates come from the same cell line and are grown and treated in reproducible ways. For higher dispersion experiments, e.g. comparing data sets from different individuals, even higher sequencing depth is required. In any case, more replicates are always as good or better than fewer replicates. Therefore, in general, we recommend performing as many replicates as possible. In our experience, most Hi-C libraries are not complex enough to produce more than 1 billion unique contacts, so reaching 6 billion contacts likely requires many replicates anyway. Sequencing depths exceeding 25 billion contacts would be required to achieve 90% well-powered loops, even with very low dispersion. Such sequencing depths are not currently tractable for most researchers but may become so in the near future as we discuss later.

### Hi-C Poweraid: a web application for differential Hi-C experiment design

Since there are various parameters to consider when planning a new Hi-C experiment that can quickly become complex and overwhelming, we built Hi-C Poweraid, an R shiny app to facilitate this planning (http://phanstiel-lab.med.unc.edu/poweraid/). Hi-C Poweraid consists of 2 main tabs: tab 1 for investigating power across different sequencing depths and tab 2 for investigating power across different loop sizes. In tab 1, the user can specify a range of replicate values, a power threshold, and one or more dispersion values. A well-powered loop is defined as one with a power above the given power threshold to detect a 2-fold change in looping interactions. This tab is useful for determining the ideal total sequencing depth and number of replicates for an experiment, and to investigate how differing dispersions can affect that decision. Tab 2 provides more granular information, albeit on just one set of parameters at a time. Here, a user can make fine-tuned selections of fold change, dispersion, replicate number, and sequencing depth per replicate. The plots provided can then be used to determine power as a function of loop size, which can help inform both the planning and interpretation of differential Hi-C experiments. For both tabs, we make use of interactive plotly plots, which allow for features such as pan and zoom, region selections, trace isolation, and hover effects to glean more specific information about certain regions or points on the plot^20^.

## DISCUSSION

Due to the expense and difficulty of performing Hi-C experiments, it is critical to first perform a careful power analysis. A power analysis can also help inform the interpretation of experiments; for example, it can help scientists determine if the small number of differential loops is due to a similarity between the samples or just a lack of power. It can also help explain biases in the sets of differential loops detected (e.g. shorter loops). To address this, we performed a rigorous power analysis of differential Hi-C experiments and developed a web application to facilitate interrogation of the resulting data. As a result of this analysis, we recommend designing experiments with at least 6 billion contacts per condition, split between 2 or more replicates to achieve a power above 0.8 to detect a 2-fold change for over 50% of loops. A more ideal, albeit currently infeasible, design would include 25 billion contacts per condition to achieve a power above 0.8 to detect a 2-fold change for over 90% of loops. The analyses and web app described here can aid in the effective use of time and resources and in justifying plans, costs, and resource distribution when proposing new experiments.

However, several caveats pertain to these estimates. First, it is important to note, “Hi-C contacts” refers to reads that have passed a variety of filtering steps and that actual sequencing depths need to be even higher than the numbers quoted here in order to achieve the appropriate number of contacts. The percentage of reads that result in Hi-C contacts varies widely based on library quality, complexity, and sequencing depth and therefore cannot be easily modeled here. Second, while Hi-C Poweraid provides good general guidelines for experimental design, optimal design is difficult to pinpoint. Different experimental designs and protocols have different dispersions that can be hard to predict but have an important impact on power. For experiments performed on multiple replicates of the same cell line, low dispersion values are expected and deep sequencing of a small number (e.g. 2) of replicates is sufficient for optimal power. Experiments in which replicates represent different donors or animals are likely to exhibit higher dispersion values and more replicates may be required to reach similar power thresholds. However, exact values for dispersion based on different experimental designs is difficult to determine. When estimating a dispersion to use for Hi-C Poweraid, we advise using the dispersion from other, similar experiments in the lab, a collaborator’s lab, or from publicly available Hi-C data. If a dispersion cannot be estimated, it is recommended to use as many replicates as is feasible, as increasing replicates is likely to increase the overall power of the experiment. Third, these results pertain to Hi-C data only and it is currently unclear how these recommendations apply to other methods, such as micro-C, Hi-ChIP, ChIA pet, and capture Hi-C. Estimates are likely to be comparable for micro-C, as the experimental design and resulting data are similar; however, protocols that involve regional enrichment (e.g. Hi-ChIP, ChIA pet, capture Hi-C, etc) will require their own power analysis.

We have found that optimally-powered Hi-C experiments require far deeper sequencing than is typically performed or feasible for most labs (e.g 25 billion contacts per condition). While current sequencing costs largely inhibit such experiments, sequencing costs have decreased drastically over the past two decades, and are likely to continue decreasing^21,22^. Multiple emerging sequencing technologies have the potential to decrease sequencing costs by 60% or more^23^. As newer technologies arise and are adopted, and as sequencing costs continue to decrease, the recommendations for sequencing depth proposed here will become more affordable and attainable.

Hi-C Poweraid is a useful tool that enables accurate Hi-C power estimates without the need to generate costly preliminary datasets or conduct complex computational analyses. These estimates will help facilitate grant proposals and provide better planning for experiments, which will ultimately translate into more robust scientific results. Well-planned experiments will improve the efficiency of allocation of time and resources, allow for more accurate interpretation of results, and expedite scientific progress.

## DATA AND CODE AVAILABILITY

Hi-C data can be accessed through SRA accession PRJNA385337 (Phanstiel et al) and GEO accession GSE63525 (Rao et al)

Hi-C Poweraid is available as an R Shiny application deployed at http://phanstiel-lab.med.unc.edu/poweraid/ The R Shiny application code is available at https://github.com/sarmapar/poweraid

## ACKNOWLEDGEMENTS

We thank Erika Deoudes for graphic design and typesetting.

## METHODS

### Hi-C Subsampling, alignment, and processing

In situ Hi-C datasets for 29 samples (18 primary and 11 replicate) of GM12878 cells were downloaded as merge_ no_dups files through GEO accession GSE63525. These files were randomly subsampled to the proper proportion of the total sequencing depth required so that the unique reads add up to the desired sequencing depth. For each line in the merge_no_dup file, a random float was generated, and if the float was less than the percent of total needed, that line was included in the new subsampled file. This method was repeated for the total sequencing depths of 50 million, 100 million, 250 million, 500 million, 750 million, 1 billion, 2 billion, 3 billion, 4 billion, and 5 billion contacts. This method resulted in files with +/- up to 0.01% difference from the whole number sequencing depths.

The 29 subsampled merge_no_dup files per sequencing depth were combined into one file per sequencing depth and processed into Hi-C files using Juicer tools^24^ (v1.14.08). Looping interactions were called at 5Kb resolution with Significant Interaction Peak (SIP) caller^25^ (v1.6.1) and Juicer tools using the merged Hi-C file from the 29, non-subsampled merge_no_dup files (5,536,073,657 total contacts) with the following parameters: “-g 2.0 -t 2000 -fdr 0.05” for a total of 14,849 loops after merging at 10Kb resolution.

The un-normalized expected and observed counts for each loop in each Hi-C file were extracted using a pre-release of mariner^26^ (v. 0.1.0). Loops were then filtered for a length shorter than 2 Mb, observed counts greater than expected, and for those only located on chromosomes 1-22, resulting in approximately 14,000 loops per sequencing depth.

### Fold change and power calculations

The fold change of observed counts for various fold changes of the counts due to looping interactions for each loop was calculated using

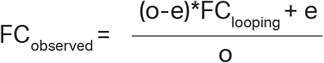

where o is observed counts, e is expected counts, and FC_looping_, is the fold change of counts due to looping interactions. For example, in **Figure 2**, the FC_looping_ values used were 2, 3, and 4.

Power was calculated per loop using the rnapower() function from the RNASeqPower package^11^, where alpha was 0.05 divided by the number of loops for a given sequencing depth, depth was the observed counts for the given loop, cv was the square root of a given dispersion, effect was the fold change of observed counts for a given fold change due to looping counts, and n was the number of given replicates. For the purpose of our analysis, dis-persion values ranging from 0.001 to 0.1, replicate values ranging from 2 to 10, and fold change values ranging from 1.1 to 10 were used.

For each combination of parameters, the percentage of well-powered loops was calculated. A well-powered loop was one where the power was greater than a set threshold, either 0.8 or 0.9 for a given fold change. A threshold of 0.8 to detect a 2-fold change was used for all analyses described here. For our initial analyses (**Fig 1-2**), we used the following parameters: 4 replicates, 2 billion reads per replicate, 0.001 dispersion.

These were chosen to reflect the same values from a previous differential Hi-C study^1^. This allowed for comparison between these 2 datasets to determine if the results from the subsampled data can be reliably extended to other Hi-C datasets and future studies (**Fig S1**).

### Visualizations

To reduce noise and to aid in visualizations, the observed and expected counts used for Figures 1 and 2 were fit to a power law curve using aomisc^27^. All figures were generated by use of the Bioconductor package plotgardener^28^.

## SUPPLEMENTAL FIGURES

**Figure S1.**
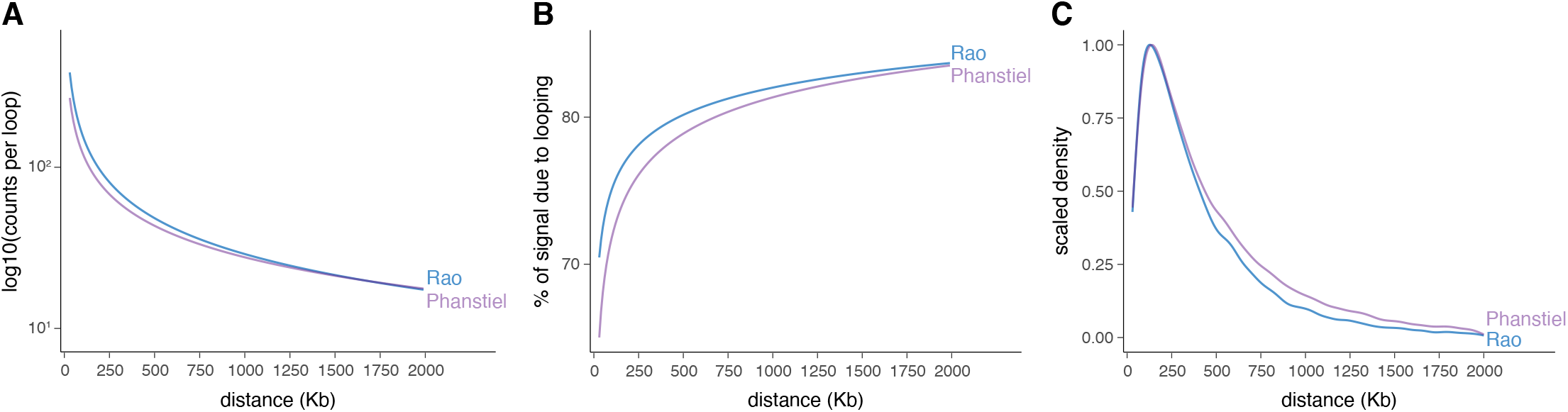
The effect of loop size on counts, loop composition, and quantity of loops from two Hi-C data sets. **(A)** Median counts vs loop size are plotted for loops from THP-1 cells (Phanstiel et al) and GM12878 cells (Rao et al). **(B)** Median percent of signal due to looping vs loop size is plotted for loops from THP-1 cells (Phanstiel et al) and GM12878 cells (Rao et al). **(C)** Distribution of loop sizes identified from THP-1 cells (Phanstiel et al) and GM12878 cells (Rao et al).

